# Identification of a distinct cluster of GDF15^high^ macrophages exhibiting anti-inflammatory activities

**DOI:** 10.1101/2024.01.11.575315

**Authors:** Chaochao Dai, Hongyu Zhang, Zhijian Zheng, Chun Guang Li, Mingyuan Ma, Haiqing Gao, Qunye Zhang, Xiaopei Cui, Fan Jiang

## Abstract

Macrophage-mediated inflammatory response may have crucial roles in the pathogenesis of a variety of human diseases. Growth differentiation factor 15 (GDF15) is a cytokine of the transforming growth factor-β superfamily, with potential anti-inflammatory activities. Previous studies observed in human lungs some macrophages which expressed a high level of GDF15. In the present study, we employed multiple techniques, including immunofluorescence, flow cytometry, and single-cell RNA sequencing, in order to further clarify the identity of such GDF15^high^ macrophages. We demonstrated that macrophages derived from human peripheral blood mononuclear cells and rat bone marrow mononuclear cells by *in vitro* differentiation with granulocyte-macrophage colony stimulating factor contained a minor population (∼1%) of GDF15^high^ cells. GDF15^high^ macrophages did not exhibit a typical M1 or M2 phenotype, but had a unique molecular signature as revealed by single-cell RNA sequencing. Functionally, GDF15^high^ macrophages were associated with reduced responsiveness to pro-inflammatory activation; furthermore, GDF15^high^ macrophages could inhibit the pro-inflammatory functions of other macrophages via a paracrine mechanism. We further confirmed that GDF15 *per se* was a key mediator of the anti-inflammatory effects of GDF15^high^ macrophage. Also, we provided direct evidence showing that GDF15^high^ macrophages were also present in other macrophage-residing human tissues in addition to the lungs. Our results suggest that the GDF15^high^ macrophage may represent a distinct cluster of macrophage cells with intrinsic anti-inflammatory functions. The (patho)physiological importance of these cells *in vivo* warrants further investigation.

## Introduction

Macrophages mediate innate immunity under homeostasis and play essential roles in regulating tissue inflammation under pathological conditions. Macrophages display remarkable heterogeneity and plasticity, and can change their physiology in response to environmental cues, giving rise to different populations with distinct functions in different organs ^1–5^. It is well established that in the body, macrophages originate from two major sources, namely embryonic hematopoietic precursors located in embryonic hematopoietic organs (such as yolk sac and fetal liver), and definitive hematopoietic stem cells located in the bone marrow ^2,6^. Most tissue-resident macrophages are derived from the embryonic origins, although to varying degrees circulating monocytes have a contribution in replenishing the pool during postnatal life. During inflammation, in contrast, the majority of macrophages infiltrating the inflamed tissue arise from circulating monocytes ^2,6^. In the lungs, for example, there are two major macrophage populations, namely alveolar macrophage and interstitial macrophage ^7,8^. With inflammatory insults, additional monocytes/macrophages are recruited from the circulation and may expand locally ^8^. Macrophage-mediated inflammation is implicated in the pathogenesis of a variety of lung diseases, including pulmonary arterial hypertension (PAH) ^7^, chronic obstructive pulmonary disease (COPD) ^9^, inflammatory acute lung injury ^10^, and pulmonary fibrosis ^11^.

Growth differentiation factor 15 (GDF15) is a cytokine of the transforming growth factor-β (TGF-β) superfamily. It was initially identified as a macrophage-derived cytokine which had inhibitory effects on lipopolysaccharide (LPS)-induced macrophage activation (thus also known as macrophage inhibitory cytokine 1/MIC-1) ^12^. In addition to macrophages, GDF15 is also highly expressed in the epithelial cells in different organs ^13^. Accumulating evidence from *in vivo* functional studies has suggested that GDF15 may be an important regulator of inflammation in the body; for example, GDF15 has been shown to have inhibitory effects on experimental systemic lupus erythematosus ^14^ and glomerulonephritis ^15^. Moreover, GDF15 exhibits protective effects in several inflammatory metabolic disorders (*e.g.* non-alcoholic fatty liver disease and type 2 diabetes mellitus) ^16^ ^17^ and cardiovascular diseases (*e.g.* myocardial infarction and atherosclerosis) ^18–20^. Traditionally, monocyte-derived macrophages are classified into M1 (pro-inflammatory) and M2 (anti-inflammatory and regenerative) phenotypes. It has been observed that GDF15 may facilitate the differentiation of macrophages toward the anti-inflammatory M2 phenotype ^20,21^. In previous studies, researchers identified some macrophage cells in human lung tissues from both healthy subjects and patients with PAH, which exhibited a high level of GDF15 expression ^22,23^. We termed these cells as GDF15^high^ macrophages thereafter in this article. GDF15^high^ macrophages were found in the alveoli; however, in PAH lungs GDF15^high^ macrophages were also present in the peri-arterial area ^22^. The functional properties of the GDF15^high^ macrophage remain unknown.

Nowadays, the cutting-edge single-cell RNA sequencing (scRNA-seq) technology empowers people to determine cell heterogeneities based on the genomic signature at a single cell level, and the emerging evidence has suggested that the macrophage heterogeneity in a single tissue is much more complex than expected ^24^. Recently, Patsalos and colleagues discovered a distinct population of macrophages with tissue regenerative functions (which were called “repair monocyte-derived macrophages”) in a murine model of sterile inflammatory muscle injury ^25^. Interestingly, these researchers demonstrated that GDF15 was not only a marker, but also a central mediator of the repair-type macrophages, which acted in local, autocrine, as well as paracrine manners to promote sustained regenerative transcriptional programs ^25^. However, the relevance of the GDF15^high^ population of macrophages in human is to be established. In the present study, therefore, we aim to clarify whether GDF15^high^ macrophages represent a functionally distinct population which has anti-inflammatory functions. We have provided evidence showing that (1) macrophages derived from human peripheral blood mononuclear cells (PBMNCs) by *in vitro* differentiation contain a minor population (∼1%) of GDF15^high^ cells; (2) scRNA-seq confirmed that GDF15^high^ macrophages represent a distinct minor population with a unique molecular signature; (3) GDF15^high^ macrophages have intrinsic anti-inflammatory functions; and (4) GDF15^high^ macrophages are present in multiple human tissues in addition to the lung.

## Materials and Methods

### Materials

Recombinant GDF15 (#10596-GD) and recombinant interferon (IFN)-γ (#585-IF) were purchased from R&D Systems (Minneapolis, MN, USA). Recombinant human granulocyte-macrophage colony stimulating factor (GM-CSF) and rat GM-CSF were from Novoprotein Technology (#C003 and #CB91 respectively) (Suzhou, Jiangsu Province, China). Monocrotaline and lipopolysaccharides (LPS) was purchased from Merck KGaA (Darmstadt, Germany).

### Human and animal ethics

Experiments involving human samples were approved by the Human Ethics Committee of Shandong University Cheeloo College of Medicine (document #KYLL-201810-054). Informed consents were obtained before commencement of the study. Experiments involving animals were approved by the Institutional Animal Ethics Committee of Shandong University Cheeloo College of Medicine (document #KYLL-2018ZM-636) and conducted in accordance with the Animals in Research: Reporting In Vivo Experiments (ARRIVE) guidelines. Animals were handled in accordance with the National Institutes of Health guide for the care and use of laboratory animals (NIH Publications No. 8023, revised 1978).

### Human tissue specimens

Human lung and kidney tissues were obtained from patients undergoing surgical tumor resections. The non-cancerous part of the tissue block was dissected and preserved for the present study. Some of the lung tissues were complicated with COPD; those without COPD were treated as normal lung tissues. Atherosclerotic arterial intima specimens were obtained from patients undergoing carotid endarterectomy. Colon tissues were either biopsies from ulcerative colitis patients undergoing colonoscopy or donations from healthy subjects undergoing routine health checks with colonoscopy. Tissue blocks were fixed in 4% paraformaldehyde solution for 24 hr at room temperature, and embedded in paraffin. Sections of 5 μm in thickness were cut.

### Isolation and differentiation of human PBMNCs

Seven healthy volunteers and 6 PAH patients were recruited in our study. Their basic demographic information was given in Supplementary Table S1. From each subject, 20 mL of peripheral venous blood was withdrawn into a heparinized tube, and diluted 1:1 in sterile PBS. PBMNCs were purified using a Ficoll-based reagent kit from TBD Science (#LTS1077, Tianjin, China). Cells (10^6^ per mL) were cultured and differentiated in RPMI 1640 medium containing 10% fetal bovine serum (all from Thermo Fisher Scientific, Waltham, MA, USA) plus 20 ng/mL of human GM-CSF as described ^26^. At day 3, half of the culture medium was replaced with fresh GM-CSF-containing medium. At day 7, adherent macrophages were collected for further experimentations.

### CD68 promoter driven-GFP transgenic reporter (CD68pro-GFP) rats

A transgenic reporter rat strain (on Sprague Dawley background) expressing enhanced GFP selectively in monocytes/macrophages was created using the strategy described previously ^27^. Briefly, the transgene construct was prepared by inserting a human CD68 promoter upstream of the GFP open reading frame. This strategy allowed stable GFP expression in both blood monocytes and mature tissue resident macrophages ^27^. The cloning service and breeding pair establishment were provided by Cyagen Biosciences (Suzhou, Jiangsu Province, China). Rats were housed in standard open-top polycarbonate cages with corn cob bedding, and maintained in an air-conditioned environment (non-SPF) with 12-hour light/dark cycles. Standard laboratory rodent diet and autoclaved tap water were provided *ad libitum*.

### Isolation and differentiation of rat bone marrow mononuclear cells (BMMNCs)

Male CD68pro-GFP rats (six weeks of age, body weight 110-120 g) were used for primary BMMNC isolation. BMMNCs were harvested from the femur and tibia as described previously ^28^. Cells were resuspended in Dulbecco’s modified Eagle medium (from Thermo Fisher) to a density of 10^6^ per mL, and cultured in the presence of 10% fetal bovine serum. Macrophage differentiation was induced by treating with rat GM-CSF at 20 ng/mL for 7 days.

### Co-culture experiments

RAW264.7 murine macrophages were cultured in Dulbecco’s modified Eagle medium with 10% fetal bovine serum as described previously ^29^. For co-culture experiments, 6-well transwell plates (0.4 μm pore size) (from Corning, New York, USA) were used. Differentiated rat macrophages were fractionated by Fluorescence-activated cell sorting (FACS). 2×10^4^ fractionated cells were resuspended in 1 mL of Dulbecco’s modified Eagle medium with 10% fetal bovine serum and seeded in the upper chamber of each well. 2×10^5^ RAW264.7 cells in the same medium were seeded in each lower chamber. Cells were co-cultured for 48 hr, then the RAW264.7 cells were either left untreated or treated with LPS (1 μg/mL) for 6 hr.

### Rat PAH models

Monocrotaline-induced PAH in CD68pro-GFP rats was established by a single bolus injection (50 mg/kg subcutaneously) as described previously ^30^.

### scRNA-seq and data processing

*In vitro* differentiated macrophages were detached with 0.25% trypsin-EDTA solution and single cell suspension was obtained by filtering through a 70-µm cell strainer. Cell viability was confirmed by staining with 0.2% trypan blue solution in PBS (pH 7.2). Library construction and double-end sequencing services were provided by BGI (Shenzhen, Guangdong Province, China), using Chromium Single Cell 3’ Reagent Kits v2 (from 10x Genomics, Pleasanton, CA, USA) and NovaSeq 6000 System (Illumina, San Diego, CA, USA) respectively. The scRNA-seq raw data were imported into Cell Ranger Single-Cell Software Suite (version 4.0, 10x Genomics) and mapped to the GRCh38 human reference genome to generate digital gene expression matrices. Subsequently, the data set was processed with Seurat R package (version 4.03), including data cleaning, doublet removal, normalization, identification of highly variable (top 2000) genes, data integration (using Canonical Correlation Analysis), and dimensionality reduction (using Principal Component Analysis) ^31^. The “Find-Neighbors” and “Find-Clusters” modules in Seurat were used for establishment of cell sub-populations, and the results were visualized using the “Run-UMAP” function. Specific marker genes were identified using the “Find-All-Markers” function with Wilcoxon rank sum test.

### Bioinformatics analysis

Differentially expressed genes (DEGs) were selected using cutoff values of fold change 1.5 and adjusted *P* 0.05 (Wilcoxon Rank-Sum test). Functional annotation and enrichment analysis for gene lists were carried out using DAVID platform (version 2021) ^32^. Cell-cell communication networks were constructed using CellChat software ^33^.

### Immunofluorescence

Anti-CD68 (#66231-2-Ig from Proteintech, Wuhan, Hubei Province, China) and anti-GDF15 (#A0185 from ABclonal, Wuhan) were used for immunofluorescence staining. Tissue sections were deparaffinized and heated in 10 mM sodium citrate buffer (pH 6.0) in a microwave oven for 15 min for antigen retrieval. Tissues were permeabilized by incubating in 0.1% Triton X-100 solution at room temperature for 10 min, and blocked in 1% bovine serum albumin solution at room temperature for 30 min. Primary antibody treatment was carried out at 4°C for overnight, followed by fluorophore-conjugated secondary antibody treatment at room temperature for 60 min. For cellular experiments, cells were cultured on glass slides. Specimens were fixed in 4% paraformaldehyde for 10 min. Other steps were the same as described above. The nuclei were counterstained with DAPI for 1 min. Fluorescent images were taken using a fluorescence microscope (Model BX43, OLYMPUS, Tokyo, Japan) and analyzed using Image-J software (NIH). For each slide, 3 - 5 random fields were selected for analysis. Morphological analysis was carried out in a blind manner.

### Flow cytometry and FACS

The following antibodies were used: anti-CD68 (#66231-2-Ig), anti-CD86 (#APC-65165), anti-CD80 (#PE-65083), anti-CD206 (#APC-65155) and anti-CD163 (#APC-65169) were from Proteintech; anti-GDF15 (#66231-2-Ig) was from ABclonal; anti-GDF15 (#BM5690) was from Boster (Wuhan); anti-TNFSF9 (tumor necrosis factor superfamily member 9) (#bs-3851R-APC) was from Bioss (Beijing, China); anti-interleukin (IL)-1β (#12-7018-41) was from Thermo Fisher; anti-IL-4 (#500808) was from BioLegend (San Diego, CA, USA). Cells detached from culture plates were filtered through 70-μm cell strainers. Rat cells were blocked with purified mouse anti-rat FcγII receptor (#550270 from BD Biosciences, Franklin Lakes, NJ, USA). Human cells were blocked by Human TruStain FcX Fc receptor blocking solution (#422301 from BioLegend). Flow cytometry analysis was performed using an AccuriC6 Plus platform; FACS was performed using a FACSAria III cell sorter (both from BD Biosciences). Data analysis and plotting were performed using FlowJo software (Version 10) (from BD).

### Quantitative real-time PCR (qPCR)

Total RNA was extracted using TRIzol Reagent (Thermo Fisher) and the concentration was determined using NanoDrop 2000 spectrophotometer (Thermo Fisher). cDNA was synthesized using PrimeScript RT-PCR Kit from TaKaRa (#RR037A) (Otsu, Shiga, Japan). The PCR reaction was carried out using SYBR green qPCR mix (#CW0957M, from CoWin Biosciences, Taizhou, Jiangsu Province, China) on StepOne Real-Time PCR System (Thermo Fisher). β-actin was used as the house-keeping gene. The relative level of mRNA expression was calculated using the 2^-ΔΔCt^ method. Sequences of the primers used were given in Supplementary Table S2.

### ELISA assay

Measurements of the protein levels of TNFSF9, GDF15 and annexin A1 were performed using ELISA kits from JYM-BIO (Wuhan). Standard curves were constructed according to the manuals; all r^2^ values of the standard curves were > 0.98.

### Data and statistics

Data were expressed as mean ± standard error of the mean (SEM). Statistical analysis was performed using GraphPad Prism (version 8.0) (from GraphPad, San Diego, CA, USA). Unpaired *t*-test was used to compare two groups; one-way analysis of variance (ANOVA) followed by *post hoc* Tukey’s test was used to compare multiple groups. All tests were run as two-tailed. A value of *P <* 0.05 was considered as significant. The *n* number shown represented independent experiments/specimens, not technical replicates.

## Results

### Existence of GDF15^high^ macrophages in the lung tissue

To precisely confirm the existence of GDF15^high^ macrophages *in vivo*, we performed immunofluorescence double labeling for CD68 and GDF15 in human lung tissues without and with COPD respectively. A small population of CD68^+^ cells with a high level of GDF15 expression could be observed in normal lung tissues (Figure 1A). Consistent with previous reports ^22^, these cells were large and round in morphology, suggesting that they were alveolar macrophages. Of note, not all of the CD68^+^ cells were GDF15^high^, indicating that GDF15^high^ macrophages constituted only a subgroup of the alveolar macrophage. GDF15^high^ macrophages were also present in COPD lungs (see Figure 1B). The abundance of GDF15^high^ macrophages in Subjects #2 and #3 was comparable to that in healthy controls; however, in Subjects #1 and #4, GDF15^high^ macrophages were sparse (Figure 1B). To delineate whether the variable pattern of GDF15^high^ macrophage distribution in COPD lungs was related to specific pulmonary pathology, we performed conventional histopathology examinations. As shown in Supplementary Figure S1, Subjects #1 and #4 had severe emphysema, resulting in poor cellularity of the lung tissue; in comparison, the lungs of Subjects #2 and #3 had significant inflammatory infiltrations. Based on these observations, we argued that the variable pattern of GDF15^high^ macrophage distribution in COPD lungs might be correlated with the degree of tissue cellularity and/or inflammatory infiltration. Corroborating the above findings, we demonstrated that GDF15^high^ macrophages were also present in rat lung tissues (Figure 1C). Consistent with the findings in human tissues, most of the rat GDF15^high^ macrophages had a morphology reminiscent of alveolar macrophage, while GDF15^low/-^ alveolar macrophages were also present throughout the tissue. The abundance of GDF15^high^ macrophages was significantly increased in the lungs from PAH models (Figure 1C). This finding was consistent with those from human samples, showing that the abundance of GDF15^high^ macrophages was significantly increased in the lungs with systemic sclerosis-associated PAH ^23^. Moreover, we observed that some of the GDF15^high^ macrophages in rat lungs (especially in PAH) were located in the interstitial area and were smaller in size, suggesting that these might be interstitial macrophages.

**Figure 1.**
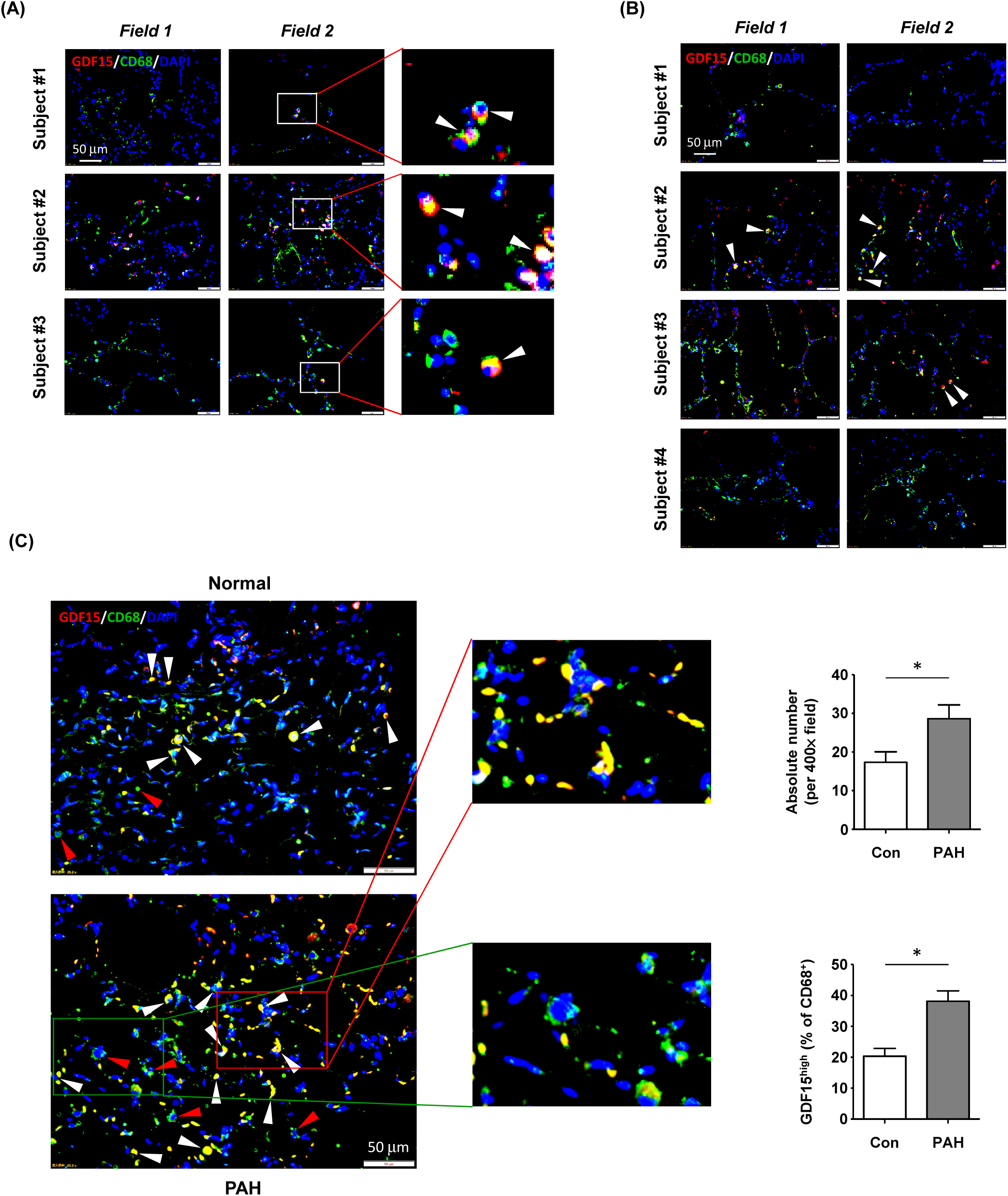
Immunofluorescence double labeling showing the existence of GDF15^high^ macrophages in lung tissues. (A) Results obtained in healthy human lung tissues from 3 independent subjects. Two representative microscopic fields were shown. Arrowheads indicated the CD68^+^GDF15^high^ macrophages. (B) Results obtained in human lung tissues with COPD from 4 independent subjects. Arrowheads indicated the CD68^+^GDF15^high^ macrophages. (C) Results obtained in rat lung tissues without and with experimental PAH. The nuclei were counterstained with DAPI (blue). White arrowheads indicated CD68^+^GDF15^high^ macrophages. The red box highlighted the presence of CD68^+^GDF15^high^ macrophages (white arrowheads); the green box highlighted the presence of CD68^+^GDF15^low^ macrophages (red arrowheads). The bar graphs showed the absolute and relative abundances of CD68^+^GDF15^high^ macrophages in normal and PAH lungs. Data were expressed as mean ± SEM. * *P <* 0.05, unpaired *t*-test, *n* = 6.

### GDF15^high^ macrophages can be differentiated from PBMNCs and BMMNCs

*In vitro* differentiation of human PBMNCs with GM-CSF yielded a population of CD68^+^ macrophages confirmed by both immunofluorescence and flow cytometry (Figure 2A and 2B). Within these induced macrophages, around 1% of them exhibited a high level of GDF15 expression as determined by flow cytometry (Figure 2C). We further confirmed the existence of such GDF15^high^ macrophages using immunofluorescence (Figure 2D). Similar results were also obtained in rat cells. Using CD68pro-GFP rats, we showed that ∼90% of GM-CSF-differentiated BMMNCs were macrophages (*i.e.* GFP expressing) (Figure 2E and 2F). As a negative control, macrophages derived from normal non-transgenic rats showed no GFP fluorescence (Figure 2E). Consistently, we demonstrated that around 1% of the BMMNC- derived macrophages were GDF15^high^ (Figure 2G and 2H).

**Figure 2.**
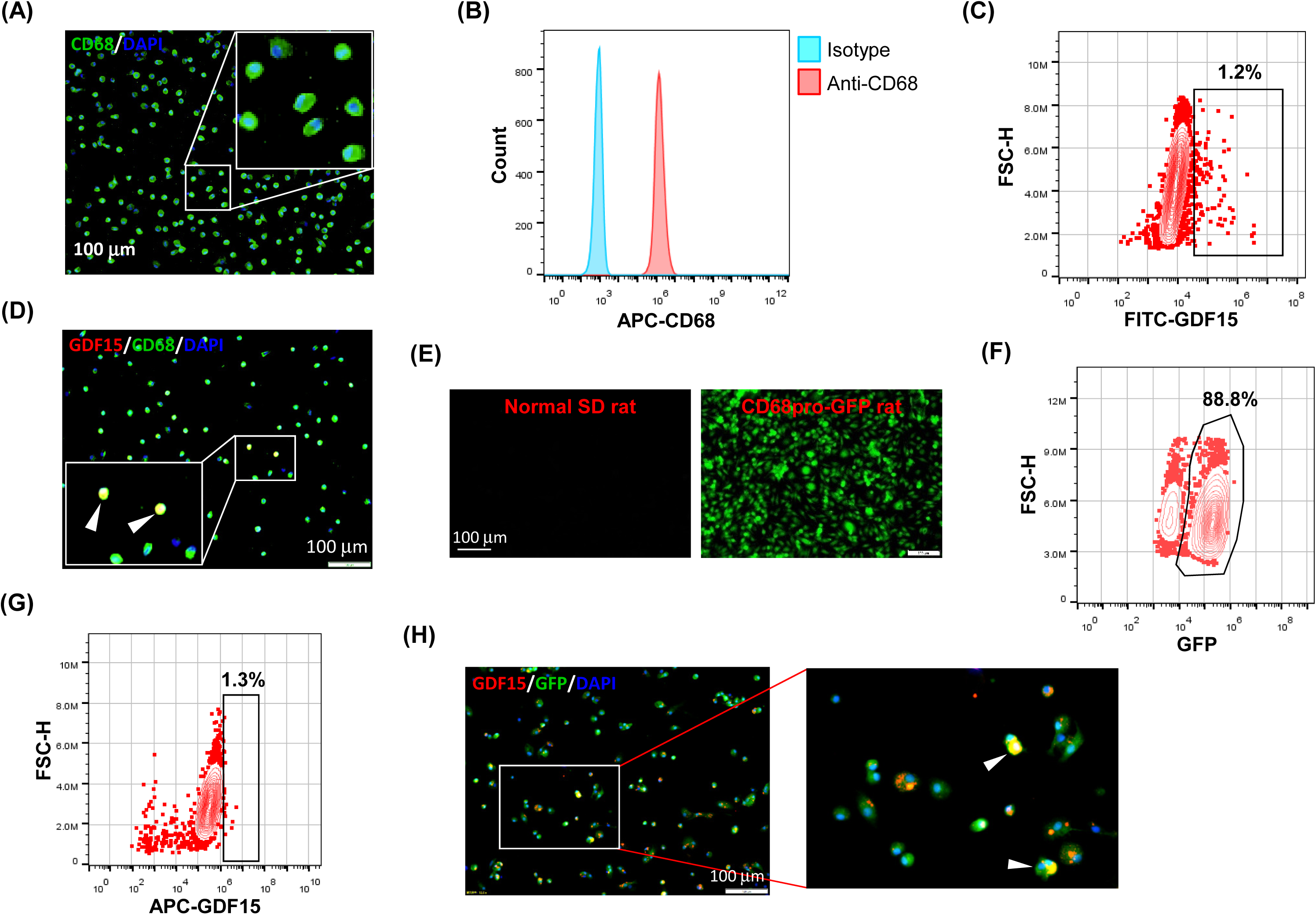
GDF15^high^ macrophages could be derived by *in vitro* differentiation of mononuclear cells. (A and B) Immunofluorescence staining and flow cytometry results confirming that *in vitro* differentiation of human peripheral blood mononuclear cells (PBMNCs) with GM-CSF for 7 days yielded CD68^+^ macrophages. (C and D) Flow cytometry and immunofluorescence double labeling results showing that the PBMNC-derived macrophages contained a minor population of GDF15^high^ cells (arrowheads in D) (example from 3 independent experiments). (E and F) Fluorescence microscopy and flow cytometry data showing that GM-CSF differentiation of rat bone marrow mononuclear cells (BMMNCs) in vitro yielded macrophages (GFP expressing) of a high purity (∼90%). CD68pro-GFP rats had a GFP transgene under the control of CD68 promoter. Cells from normal rats showing no GFP fluorescence served as a negative control (left panel in E). (G and H) Flow cytometry and immunofluorescence double labeling data (from 3 independent experiments) showing that the BMMNC-derived macrophages (from CD68pro-GFP rats) contained a minor population of GDF15^high^ cells (arrowheads in H). The flow cytometry data in panels C and G were from cells gated for GFP^+^. The nuclei were counterstained with DAPI (blue).

### GDF15^high^ macrophages do not exhibit a typical M1 or M2 phenotype

To understand whether the GDF15^high^ macrophages belonged to the conventional M1 or M2 phenotype, we performed flow cytometry characterization in human PBMNC-derived macrophages. We tested the M1 markers CD86, CD80 and IL-1β, and the M2 markers CD206, CD163 and IL-4. As shown in Figure 3, GDF15^high^ macrophages were primarily CD206^+^. CD163 and CD86 were only expressed in part of the GDF15^high^ macrophages (*i.e.* CD163^+/-^ and CD86^+/-^). However, we found that there were nice correlations between the expression levels of GDF15 and those of CD80, IL-1β and IL-4 (Figure 3). Taken together, these data indicate that the GDF15^high^ population belongs to neither the conventional M1 phenotype nor the M2 phenotype.

**Figure 3.**
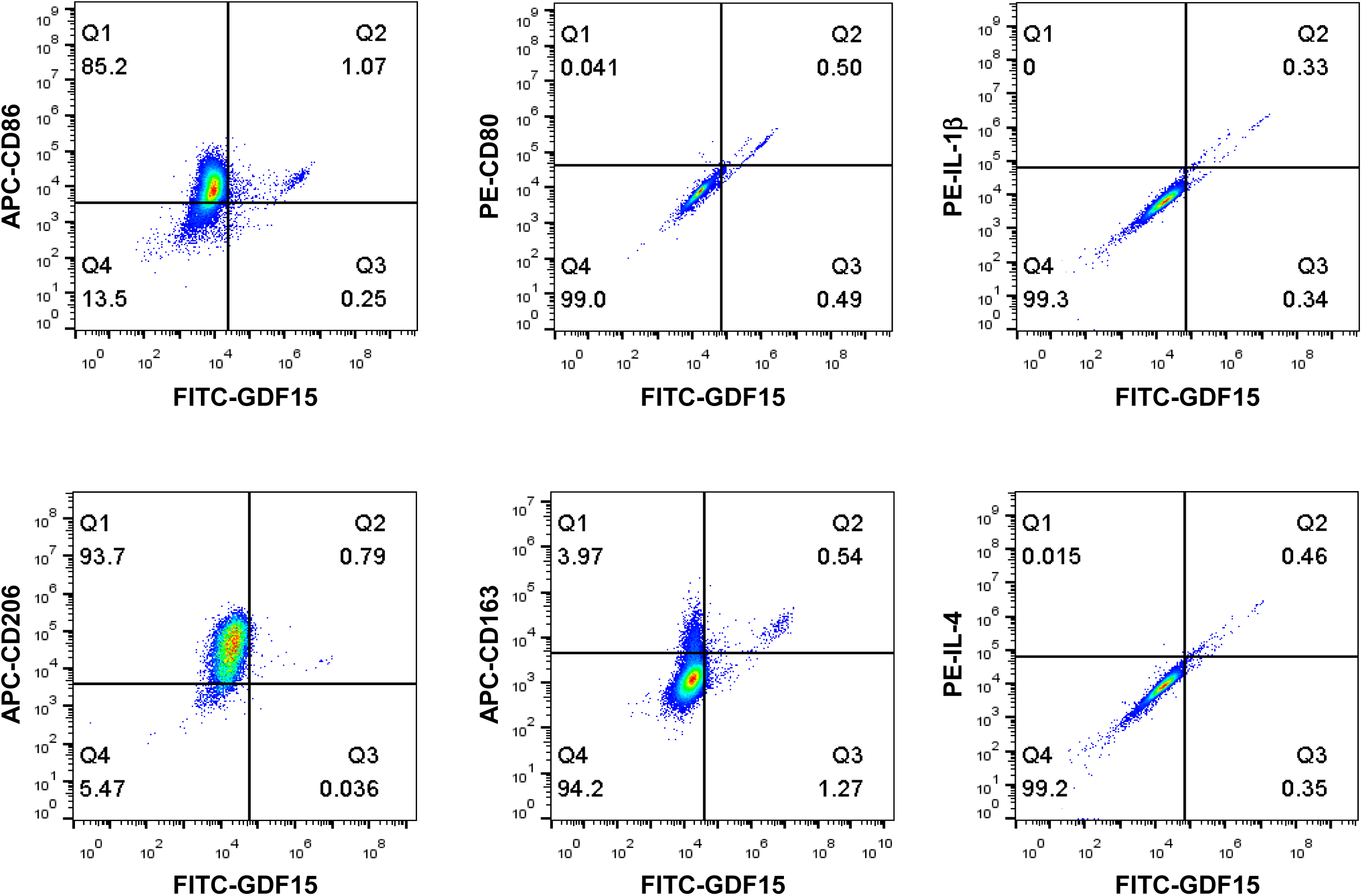
Flow cytometry results showing that GDF15^high^ macrophages did not exhibit a typical M1 or M2 phenotype. Experiments were performed in human PBMNC-derived macrophages, using CD86, CD80 and IL-1β as the M1 markers, and CD206, CD163 and IL-4 as the M2 markers. Data were from a single test using pooled samples from 4 healthy volunteers.

### Molecular characterization of GDF15^high^ macrophages using scRNA-seq

To further confirm the existence of GDF15^high^ macrophages and delineate the molecular characteristics of these cells, we applied scRNA-seq in human PBMNC-derived macrophages. We obtained PBMNC-derived macrophages from 3 healthy volunteers, 3 PAH patients harboring mutations in bone morphogenetic protein receptor type II (BMPR2), and 3 PAH patients without BMPR2 mutations. Combined analysis of all 9 samples by projecting to global uniform manifold approximation and projection (UMAP) plots led to the identification of 7 distinct macrophage sub-populations (Figure 4A and 4B). Separate analysis and comparison of the data between healthy controls and PAH patients showed virtually identical clustering profiles of the macrophages (Figure 4B inset). Moreover, quantitative analysis of the relative abundance of the macrophage sub-populations demonstrated that there were no significant differences between healthy subjects and PAH patients with or without BMPR2 mutations (Supplementary Figure S2). The putative nomenclatures for these macrophage sub-populations were given in Figure 4B. The expression patterns of identified marker genes for the different macrophage sub-populations were shown in Figure 4C. Of note, our scRNA-seq data confirmed the existence of a cluster of macrophages which exhibited a relatively high level of GDF15 expression (*i.e.* cluster 5 in Figure 4B).

**Figure 4.**
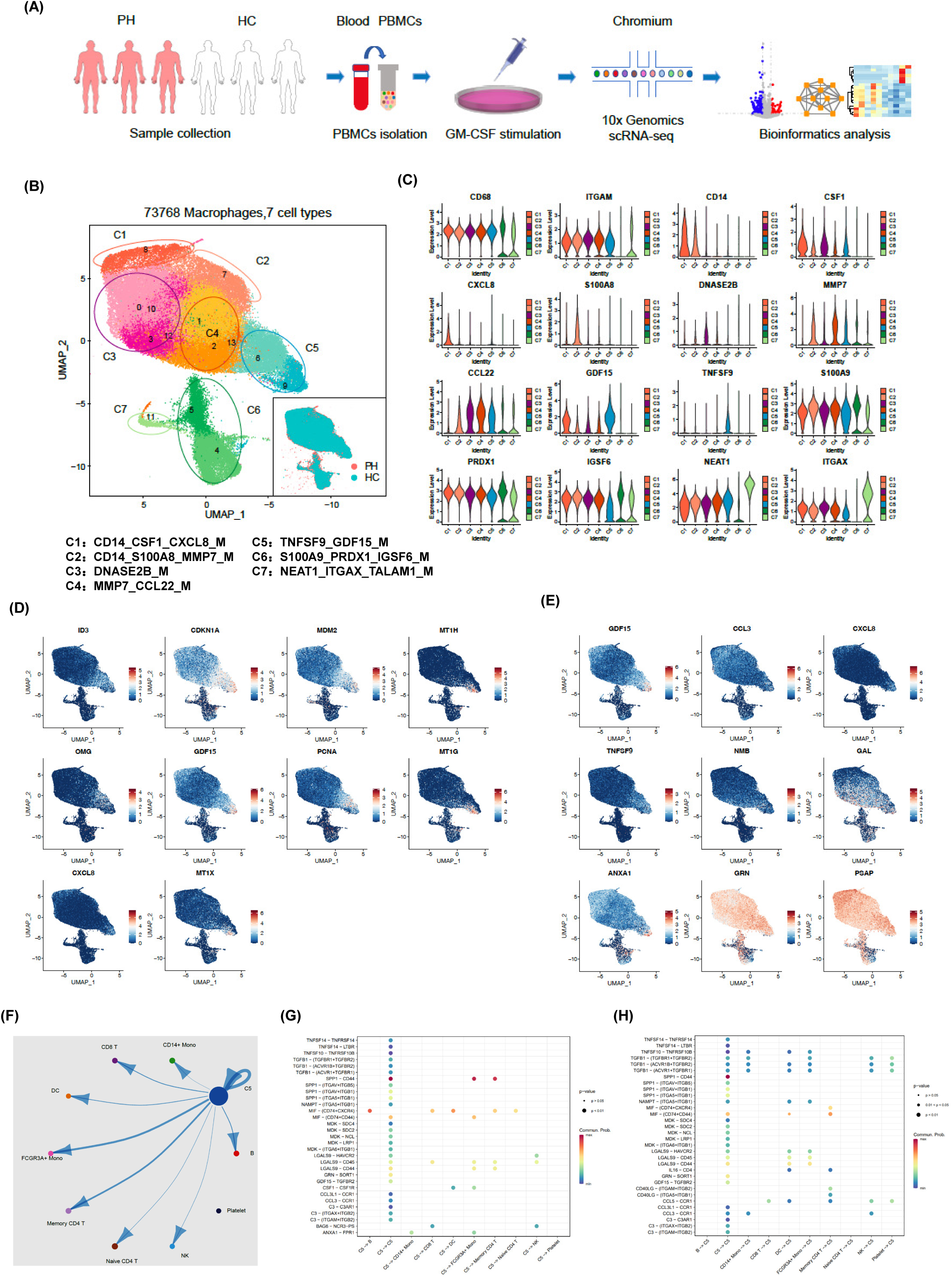
Molecular characterization of human PBMNC-derived GDF15^high^ macrophages with scRNA-seq. (A) Graphical outline of the experimental procedure. (B) UMAP plots showing the identified cell clusters (C1 to C7) based on the scRNA-seq data from total 73,768 cells combined from samples from 3 healthy volunteers, 3 PAH patients harboring mutations in BMPR2 gene, and 3 PAH patients without BMPR2 mutations. The inset showed that macrophages from healthy controls and PAH patients had virtually identical clustering profiles when analyzed separately. The putative nomenclatures for C1 to C7 were given below the graph. (C) Violin plots showing the expression patterns of identified marker genes for C1 to C7. (D) UMAP plots showing expression patterns of the top 10 genes that were overexpressed in GDF15^high^ macrophages (C5) as compared to GDF15^low^ cells. (E) UMAP plots showing expression patterns of the top 9 genes encoding secreted proteins which were overexpressed in GDF15^high^ macrophages as compared to GDF15^low^ cells. (F) Cell-cell communication network map created using CellChat showing the possible effector cells of the GDF15^high^ macrophage. (G and H) Predicted ligand-receptor pairs potentially involved in the signaling of reciprocal communications between GDF15^high^ macrophage and other cell types as listed in panel F.

Next, we searched for DEGs between GDF15^high^ and GDF15^low^ macrophages and performed functional clustering analysis. In cells from healthy controls, 81 genes were significantly upregulated in GDF15^high^ macrophages, while 58 genes were downregulated (Supplementary Figure S3 and Supplementary Table S3). The total number of DEGs in cells from PAH patients was less than that of healthy subjects; however, these regulated genes were highly consistent with those found in healthy subjects (see Supplementary Figure S3). The top ten DEGs and top nine DEGs encoding secreted proteins were shown in Figure 4D and 4E. To validate the above genomics data, we performed ELISA for TNFSF9, GDF15 and annexin A1 (ANXA1) as examples in the conditioned medium of rat BMMNC-derived macrophages. We found that the expressions of these genes were all upregulated in GDF15^high^ cells (Supplementary Figure S4). Functional clustering analysis on the DEGs revealed that GDF15^high^ macrophages were associated with a transcriptional signature featuring altered expressions of genes involved in cellular redox homeostasis and detoxification. These functional clusters included GO (Gene Ontology) terms cellular response to cadmium ion, cellular response to metal ion, detoxification of copper ion, cellular response to zinc ion, and cellular response to copper ion; UniprotKB Keywords cadmium, metal-thiolate cluster, and copper; InterPro classifications metallothionein domain, metallothionein superfamily (eukaryotic), and metallothionein (vertebrate, metal binding site) (all Benjamini *P* < 0.05). Specific genes in this regard included upregulated expressions of metallothioneins (MT1E/1G/1H/1M/1X/2A) and ferredoxin reductase (FDXR); and downregulated expressions of thioredoxin interacting protein (TXNIP) (a pro-oxidant factor), antioxidant 1 copper chaperone (ATOX1) and ferredoxin 1 (FDX1). Overall, these changes indicated possibly an increased redox defensive/detoxification capability in GDF15^high^ macrophages. We also paid attention to genes which were known to be crucial for fundamental macrophage functions. Among the DEGs, there were downregulations in cathepsins (CTSB/D/Z), macrophage scavenger receptor 1 (MSR1), major histocompatibility complexes (HLA-DQB1 and HLA-DRB1), Fc gamma receptor Ia (FCGR1A), Fc epsilon receptor Ig (FCER1G), and the reactive oxygen species-producing NADPH oxidase subunit cytochrome b-245 beta chain (CYBB, aka NOX2 or gp91phox). Moreover, we noticed that, in addition to GDF15, several genes with documented anti-inflammatory functions were upregulated, including annexin A1, dual specificity phosphatase 14 (DUSP14), and TNF alpha induced protein 6 (TNFAIP6). These bioinformatics results indicate that GDF15^high^ macrophage might have distinct functional properties as compared to the conventional macrophage.

Our initial UMAP analysis revealed that the GDF15^high^ macrophage comprised two sub-clusters (see the different colors shown in Figure 4B), which located adjacent to each other on the UMAP plot. Attempting to uncover the interrelationship between the two sub-clusters, we performed RNA velocity analysis ^34^. As shown in Supplementary Figure S5, these sub-clusters represented cells of the same identity but with diverging differentiation potentials; it was unlikely that these sub-clusters denoted two sequential differentiation status on the same differentiation route.

To identify possible effector cells of the GDF15^high^ macrophage, we created a cell-cell communication network map using CellChat software, which utilized a mass action-based model to quantify the probability of cell-cell communications on the basis of known ligand-receptor pairs and the transcriptome profiles of sequenced cells ^33^. The input data sets were from the GDF15^high^ macrophage population and the total peripheral blood mononuclear cell pool. As shown in Figure 4F, the most significant target cell types included memory CD4^+^ T cell, non-conventional FCGR3A^+^ monocyte, and the GDF15^high^ macrophage *per se*. The ligand-receptor pairs potentially involved in reciprocal communications between GDF15^high^ macrophage and other cell types were shown in Figure 4G and 4H. These ligand-receptor pairs included signaling pathways with recognized anti-inflammatory functions, such as TGF-β signaling, colony stimulating factor 1 (CSF1) signaling, and annexin A1 signaling. It was noted that GDF15 signaling appeared to be implicated only in macrophage-macrophage interactions.

Finally, to understand whether there was possibly a special sub-population of GDF15- expressing monocytes which could give rise to mature GDF15^high^ macrophage, we utilized a public scRNA-seq database, which integrated 178,651 mononuclear phagocytes encompass dendritic cells, monocytes, and macrophages from 13 different tissues ^35^ (the interface available at https://macroverse.gustaveroussy.fr/2021_MoMac_VERSE), and found that there was not a discernable cluster of monocytes with enhanced expression of GDF15 (Supplementary Figure S6), suggesting that the GDF15^high^ sub-population was formed *de novo* during the process of monocyte to macrophage differentiation.

### Inflammatory activation is dampened in GDF15^high^ macrophages

Because GDF15 is mainly expressed intracellularly, we searched substitute cell surface markers that were suitable for FACS purification. By reanalyzing DEGs in the GDF15^high^ population using the scRNA-seq data, we identified several potential marker genes including OMG (oligodendrocyte myelin glycoprotein), TNFSF9 (tumor necrosis factor superfamily member 9), FAS (Fas cell surface death receptor) and PKD2L1 (polycystin 2 like 1 transient receptor potential cation channel) (Figure 5A). Since TNFSF9 (also known as 4-1BB ligand and CD137 ligand) is a typical transmembrane glycoprotein readily expressed in macrophages ^36^, we examined whether TNFSF9 could be a substitute marker for GDF15 using flow cytometry. We verified that rat BMMNC-derived macrophages contained a minor fraction of TNFSF9^high^ cells, whose expression level was correlated with that of GDF15 (Figure 5B). We also confirmed the correlated expression patterns between TNFSF9 and GDF15 in human PBMNC-derived macrophages (Figure 5C). Using TNFSF9 as a substitute marker, we obtained GDF15^high^ and GDF15^low^ macrophages (rat BMMNC-derived), which were then primed with IFN-γ and further stimulated with LPS. We demonstrated that GDF15^high^ macrophages displayed dampened inflammatory reactions to LPS stimulation as compared to GDF15^low^ macrophages, as evidenced by the lowered expressions of TNF-α, IL-1β and IL-6 (Figure 5D), reduced migratory activity (Figure 5E), and reduced phagocytosis (Figure 5F). GDF15^high^ macrophages showed decreased cell migration even in the absence of LPS stimulation.

**Figure 5.**
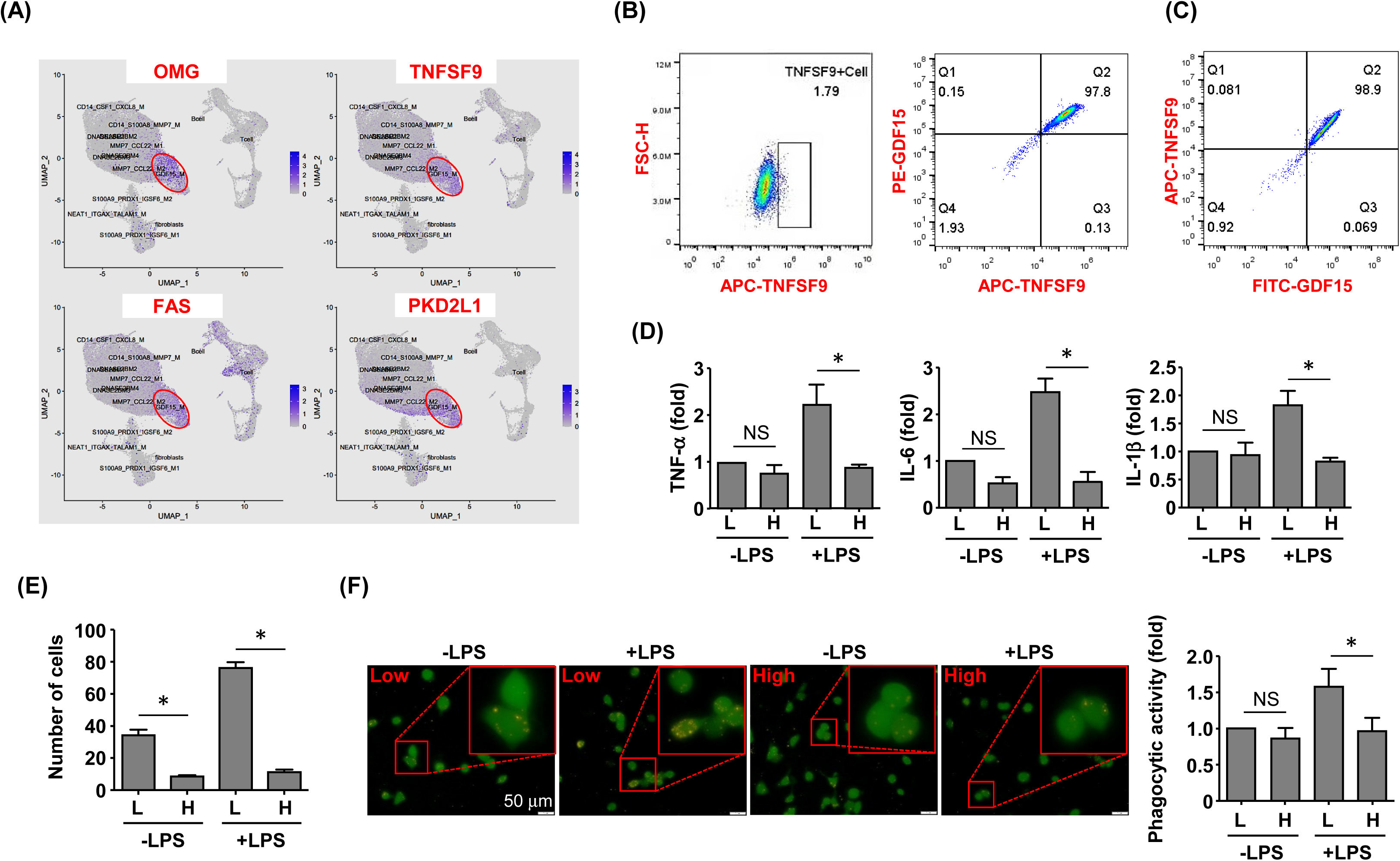
GDF15^high^ macrophages exhibited reduced inflammatory activation in vitro. (A) Expression patterns of potential substitute cell surface markers for GDF15 based on the scRNA-seq data. (B) Flow cytometry results showing that rat BMMNC-derived macrophages contained a minor fraction of TNFSF9^high^ cells, whose expression level was correlated with that of GDF15 (from 3 independent experiments). (C) Flow cytometry verification of the correlation between TNFSF9 and GDF15 expressions in human PBMNC-derived macrophages (from 2 independent experiments). (D) Real-time PCR results showing that GDF15^high^ macrophages (H) exhibited reduced expressions of TNF-α, IL-1β and IL-6 in response to LPS stimulation (1 μg/mL for 6 hr), as compared to GDF15^low^ cells (L). Rat BMMNC-derived macrophages were FACS purified using TNFSF9 as a substitute marker for GDF15, and primed with IFN-γ (10 ng/mL for 12 hr). (E) Boyden chamber cell migration assay showing that GDF15^high^ macrophages (H) exhibited reduced migratory activity as compared to GDF15^low^ cells (L) in the absence and presence of LPS stimulation. (F) Representative fluorescent microscopic images and quantitative data showing that GDF15^high^ macrophages exhibited reduced phagocytic activity in the presence of LPS stimulation as compared to GDF15^low^ cells. Phagocytosis was assessed by internalization of fluorochrome-labeled latex beads (orange color). The macrophages were from CD68pro-GFP rats. Data were mean ± SEM. * *P <* 0.05, one-way ANOVA, *n* = 3 in D; 6 in E; 6 in F. NS, no significance.

### GDF15^high^ macrophages can produce anti-inflammatory effects via paracrine mechanisms

To clarify whether GDF15^high^ macrophages could affect the functions of other macrophages in a pro-inflammatory environment, we co-cultured murine RAW264.7 cells with GDF15^high^ and GDF15^low^ macrophages (rat BMMNC-derived) separately for 48 hr, and then the RAW264.7 cells were further challenged with LPS. We demonstrated that RAW264.7 cells co-cultured with GDF15^high^ macrophages showed reduced cytokine expressions and reduced cell migration, while the phagocytic activity was not different (Figure 6A-6C). The inhibitory effects of GDF15^high^ macrophages were also partly observed in unchallenged RAW264.7 cells. To further validate these results, we repeated the co-culture experiments by replacing the RAW264.7 cell line with primary macrophages (derived from rat BMMNCs). As shown in Figure 6D-6F, similar anti-inflammatory effects of the GDF15^high^ macrophage were all observed in these experiments. Unlike RAW264.7, the phagocytic activities of primary macrophages were decreased in the GDF15^high^ co-culture (both in the presence and absence of LPS stimulation) (Figure 6F). In order to confirm that the decreases in inflammatory reactions in the GDF15^high^ co-culture were indeed a result of paracrine release of certain anti-inflammatory factor(s) from GDF15^high^ macrophages, we tested whether GDF15 itself could be the responsible paracrine factor as inspired by the previous studies ^12,37,38^. Firstly, we demonstrated that pretreatment with exogenous GDF15 mimicked the anti-inflammatory effects of GDF15^high^ macrophage in LPS-challenged RAW264.7 cells (including reduced cytokine expressions and reduced cell migration, but with no effect on phagocytosis) (Figure 7A-7C). Secondly, in both RAW264.7 cells and primary BMMNC-derived macrophages, we showed that the conditioned-medium from GDF15^high^ macrophages exhibited similar anti-inflammatory effects as GDF15^high^ co-cultures (Figure 7D and 7E), while co-incubation with a neutralizing antibody of GDF15 diminished the anti-inflammatory effects of the conditioned-medium.

**Figure 6.**
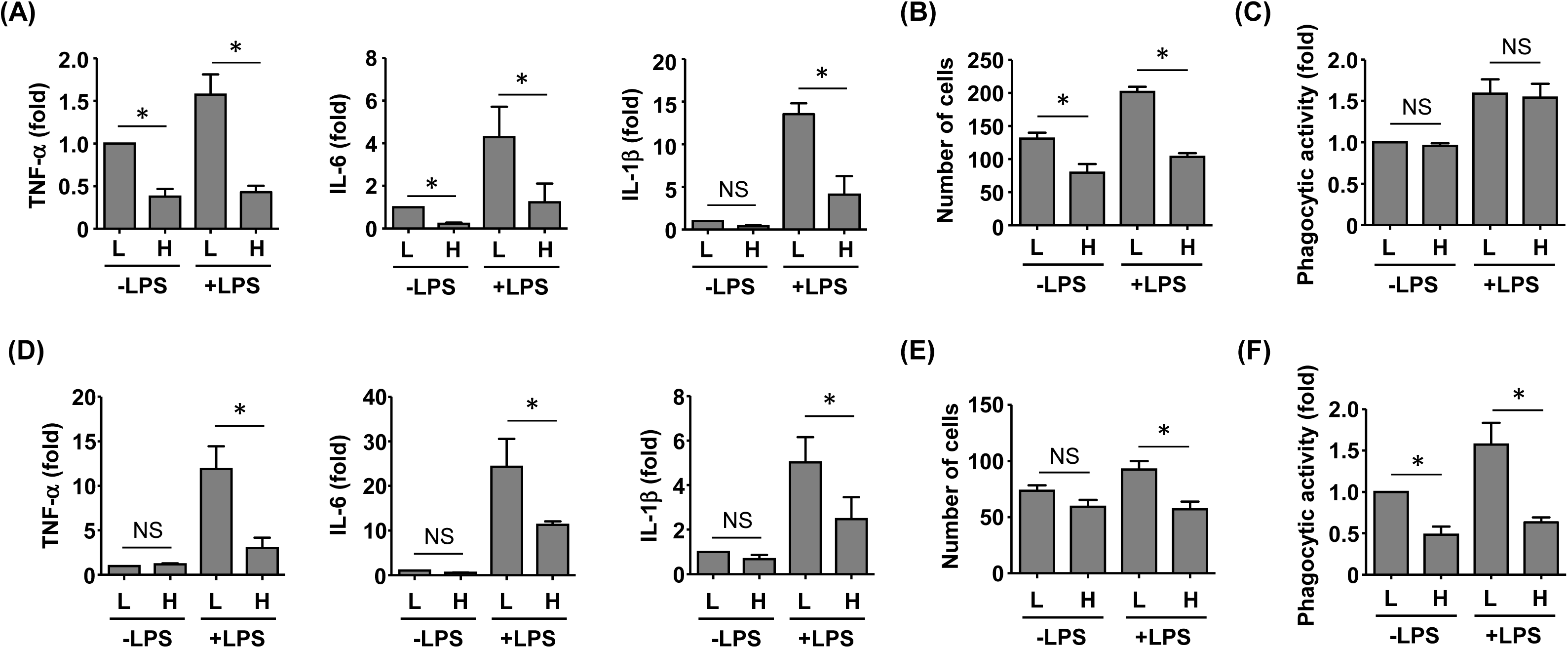
GDF15^high^ macrophages exerted anti-inflammatory effects via paracrine mechanisms. (A to C) RAW264.7 cells co-cultured with rat BMMNC-derived GDF15^high^ (H) or GDF15^low^ (L) macrophages were left untreated or stimulated with LPS for 4 hr. Results for the expression of pro-inflammatory cytokines (A, real-time PCR), cell migratory activity (B), and phagocytic activity (C) were shown. (D to F) Unsorted rat BMMNC-derived macrophages co-cultured with rat GDF15^high^ (H) or GDF15^low^ (L) macrophages were left untreated or stimulated with LPS for 4 hr. Results for the expression of pro-inflammatory cytokines (D, real-time PCR), cell migratory activity (E), and phagocytic activity (F) were shown. Data were mean ± SEM. * *P <* 0.05, one-way ANOVA, *n* = 4-5 in A; 6 in B; 3 in C; 3-4 in D; 6 in E; 6 in F. NS, no significance.

**Figure 7.**
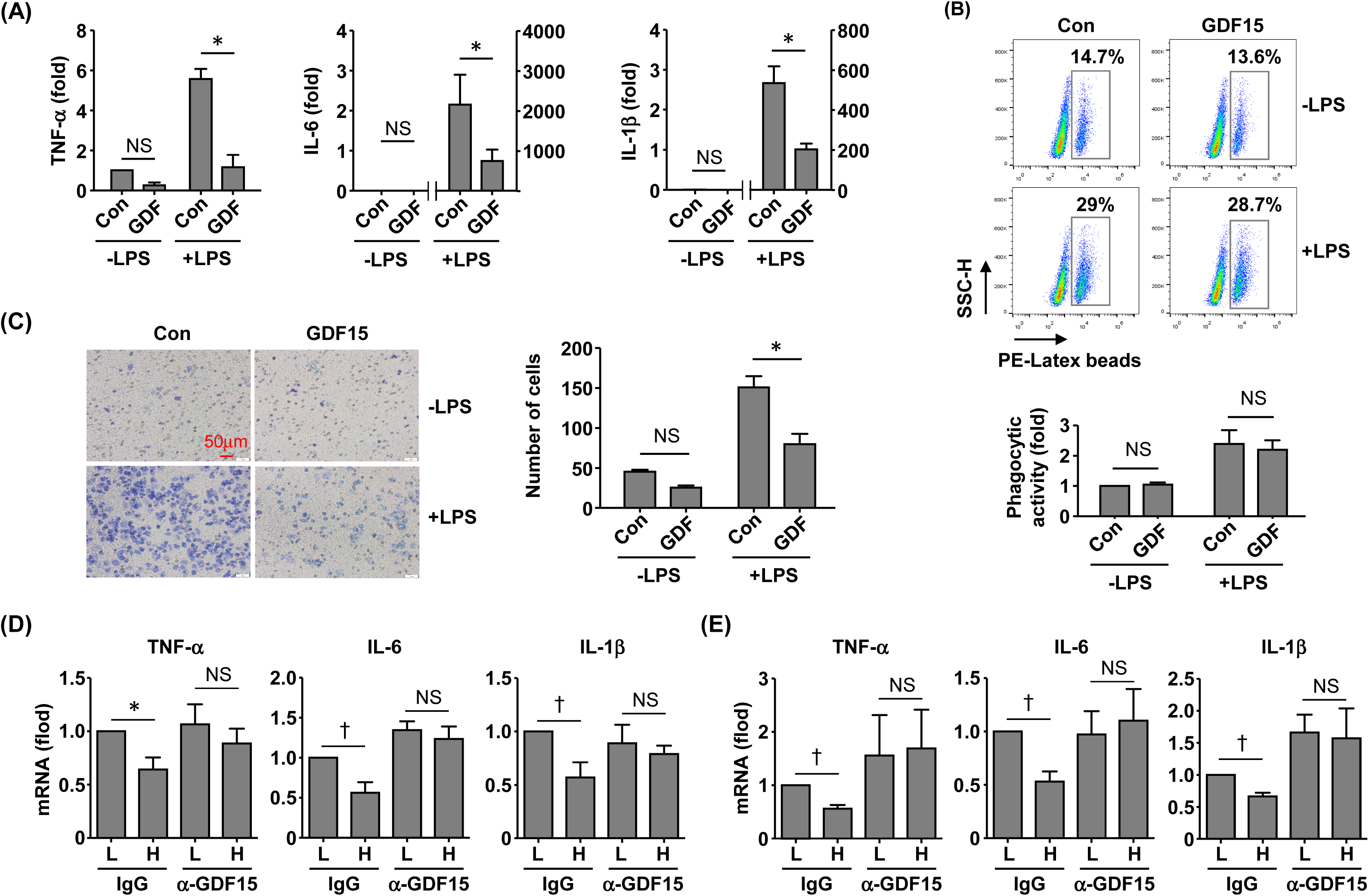
GDF15 might be a macrophage-derived anti-inflammatory factor. (A) Real-time PCR results showing that treatment with exogenous GDF15 (20 ng/mL) inhibited LPS-induced expression of pro-inflammatory cytokines in RAW264.7 cells. (B) Flow cytometry results showing that GDF15 treatment had no effects on phagocytosis in RAW264.7 cells without or with LPS stimulation. (C) Representative images and quantitative data of Boyden chamber assay showing that exogenous GDF15 inhibited migration of LPS-challenged RAW264.7 cells. Cells on the membrane were stained with Giemsa. (D) Effects of conditioned medium from GDF15^high^ macrophages (H), as compared to the medium from GDF15^low^ cells (L), on the expression of pro-inflammatory cytokines in RAW264.7 cells. All experiments were performed in the presence of LPS stimulation. α-GDF15, GDF15neutralizing antibody; IgG, non-specific immunoglobulin control. (E) The same experiments as those in D carried out in rat BMMNC-derived macrophages. Data were mean ± SEM. * *P <* 0.05, one-way ANOVA; † *P <* 0.05, unpaired *t*-test, *n* = 3-5 in A; 3 in B; 6 in C; 3 in D; 3 in E. NS, no significance.

### GDF15^high^ macrophages are present in multiple human tissues

In addition to the lungs, we also examined other human tissues which were known to contain abundant tissue-resident macrophages, including intestine ^5^ and kidney ^3^. In colon tissues from both healthy subjects and patients with ulcerative colitis, GDF15^high^ macrophages were detected in the lamina propria layer (Figure 8A). However, as compared to normal tissues, we failed to detect bulk macrophage infiltration in the colitis tissues as expected (Figure 8A); this pattern of macrophage distribution in inflammatory bowel disease was similar to the observations reported by Wright and colleagues ^39^. In parallel, there were no significant differences in the absolute or relative prevalence of GDF15^high^ macrophages between the healthy subjects and the patients with ulcerative colitis (Figure 8A). GDF15^high^ macrophages were also identified in kidney tissues (Figure 8B). Moreover, we found that GDF15^high^ macrophages were present in atherosclerotic plaques (Figure 8C), which were lesions profoundly composed of infiltrating macrophages in the sub-endothelial space ^40^. This result was consistent with previous studies ^41^.

**Figure 8.**
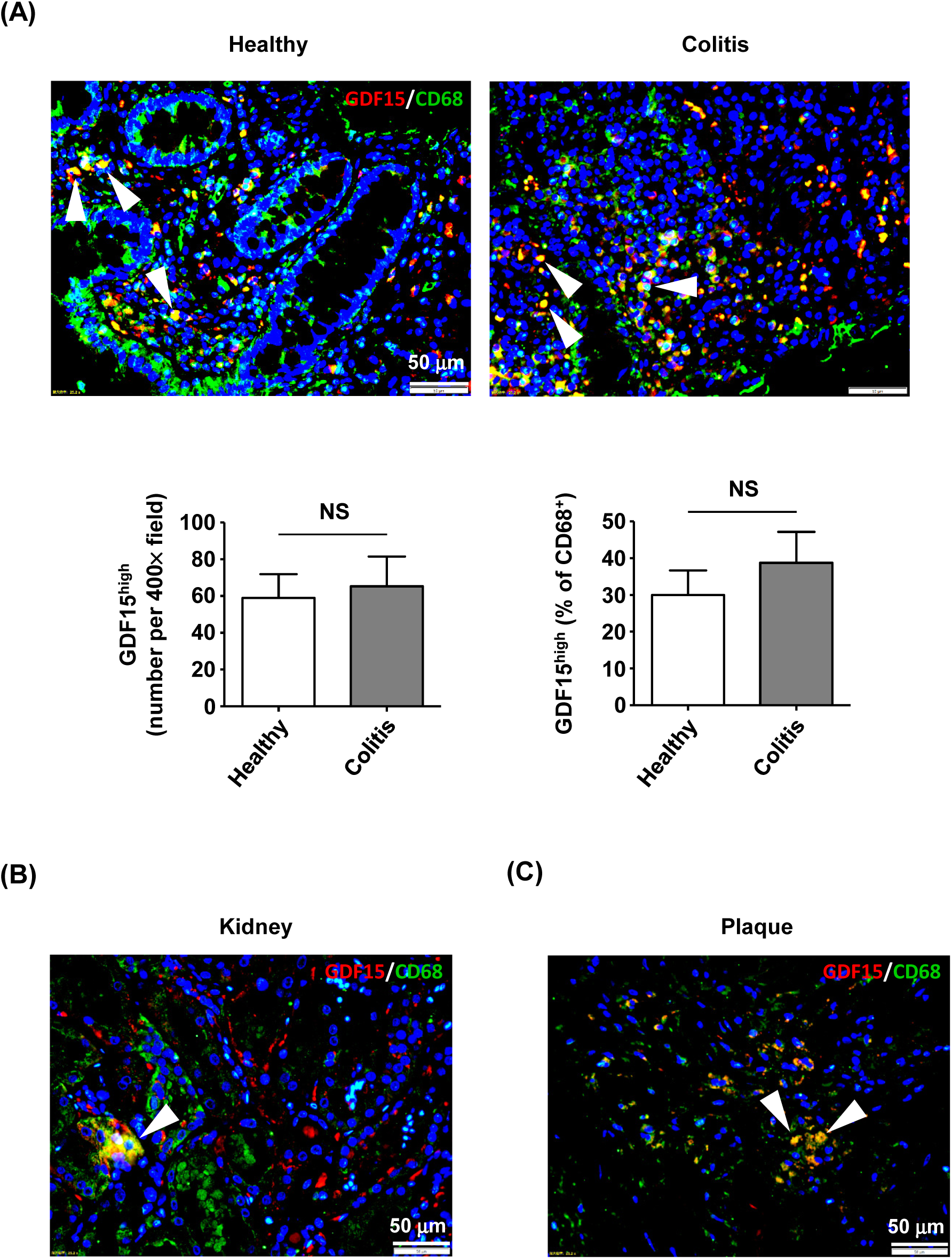
Detection of GDF15^high^ macrophages in various human tissues. GDF15^high^ macrophages (arrowheads) were identified using immunofluorescence double labeling with anti-CD68 (green color) and anti-GDF15 (red color) antibodies in (A) colon tissues from both healthy subjects and patients with ulcerative colitis, (B) kidneys (the normal peri-tumor tissue) (tested in one sample only) and (C) atherosclerotic plaques in the carotid artery (representative data from 6 independent samples showing similar results). The nuclei were counterstained with DAPI (blue). Data were mean ± SEM. NS, no significance, unpaired *t*-test, *n* = 3 for Healthy; 4 for Colitis.

## Discussion

In the present study, we have verified the existence of a minor population of macrophages expressing a high level of GDF15 (GDF15^high^) in humans and animals, which have an inhibitory effect on the pro-inflammatory functions of other macrophages via a paracrine mechanism. Our findings have confirmed the previous histopathological observations showing the existence of GDF15^high^ macrophages in human lungs ^22,23^. In addition, we have provided direct evidence that these GDF15^high^ macrophage cells are also present in other tissues/organs, at least under pathological conditions. Moreover, our results support the recent discovery of a population of monocyte-derived, GDF15-expressing macrophages in muscles, which have crucial roles in mediating tissue regeneration following inflammatory injuries ^25^. We have showed that the GDF15^high^ macrophage does not strictly belong to either the M1 or the M2 phenotype ^4^, highlighting the high degree of heterogeneity of macrophages which is more complex than that traditionally thought, as revealed by recent scRNA-seq studies ^2,24^.

The developmental origin(s) of GDF15^high^ macrophages in the body is not clear. We have shown that GDF15^high^ macrophages can be derived from peripheral blood mononuclear cells by *in vitro* differentiation. Consistently, Patsalos *et al.* provided evidence suggesting that the GDF15-expressing regenerative macrophages found in injured muscles were infiltrating monocyte-derived in nature. Moreover, we have found that there are more GDF15^high^ macrophages in the lungs from experimental PAH models, where the inflammatory macrophage cells appear to mainly originate from blood-borne monocytes ^42^. Taken together, these observations suggest that GDF15^high^ macrophages may accrue locally as a consequence of monocyte infiltration at least during inflammatory insults. On the other hand, GDF15^high^ macrophages are also present in non-inflamed normal lungs, indicating that these cells are part of the tissue-resident macrophages. However, whether this GDF15^high^ population has an embryonic origin remains to be clarified.

Our results indicate that GDF15^high^ macrophages are associated with dampened responsiveness to pro-inflammatory activation; furthermore, they also inhibit the pro-inflammatory functions of other macrophages via a paracrine mechanism, and GDF15 *per se* appears to be a key mediator of the anti-inflammatory effects of GDF15^high^ macrophages. These findings are supported by the study of Patsalos *et al.*, showing that the GDF15- expressing regenerative macrophages have anti-inflammatory properties ^25^. Specifically, GDF15 gene deletion resulted in an increase in the accumulation of F4/80^+^ macrophages in injured muscles, while exogenous GDF15 administration decreased the total number of infiltrating CD45^+^ myeloid cells ^25^. In Kupffer cells, Li *et al.* have demonstrated that GDF15 treatment attenuated LPS-induced expressions of inflammatory cytokines and inducible nitric oxide synthase ^38^. Interestingly, GDF15, being a paracrine anti-inflammatory factor, has also been observed in rat cardiac progenitor cells, of which allogeneic transplantation in the myocardium represses T cell-mediated inflammation and promotes M2 macrophage polarization ^20^. In addition, GDF-15 was also shown to inhibit the recruitment of polymorphonuclear cells following ischemic tissue injuries ^19^. Currently, the signaling mechanisms underlying the anti-inflammatory activity of GDF15 are not clearly understood. The (epi)genetic mechanisms governing the differentiation of GDF15^high^ macrophages remain elusive. Our present results suggest that GDF15^high^ macrophages are formed *de novo* during macrophage differentiation and/or maturation, but do not support the existence of a population of dedicated precursor cells (*i.e.* GDF15^high^ monocytes).

The concept of “regulatory macrophage” was proposed more than a decade ago, based on the observations that some macrophages could produce a high level of IL-10 and exert inhibitory effects on immune responses ^1^. However, the identity of such a cluster of anti-inflammatory macrophages remains poorly defined because the current markers used to distinguish these cells are highly inconsistent and even controversial ^43^, and the borderline between regulatory macrophage and M2 macrophage is vague ^44^. The findings from the present study and other researchers ^25^ raise several interesting questions: (1) Can the functions of GDF15^high^ macrophage fulfill the requirement for a regulatory macrophage? (2) What are the specific surface markers that can be used to precisely define this macrophage sub-population? (3) What is the secretome profile of GDF15^high^ macrophage that is associated with its anti-inflammatory and tissue regenerative functions? Our study has confirmed that the distribution of GDF15^high^ macrophages is wide-spread in the body, at least under pathological conditions, indicating that these cells might be associated with some generic (rather than tissue-restricted) biological functions. In addition to the lungs, the presence of GDF15^high^ macrophages has also been reported in human atherosclerotic plaques ^41^. Moreover, our data suggest that during sterile inflammation, there is a trend of increase in the abundance of GDF15^high^ macrophages in the inflamed tissue, a process that might represent an intrinsic feedback mechanism to facilitate inflammation resolution and subsequent tissue repair. Nonetheless, the physiological and pathophysiological significance of GDF15^high^ macrophages *in vivo* is to be clarified by additional studies.

In summary, we have demonstrated that a minor population of peripheral and bone marrow mononuclear cells can be differentiated *in vitro* into macrophages that express a high level of GDF15. These GDF15^high^ macrophages do not exhibit a typical M1 or M2 phenotype but are associated with a unique molecular signature as revealed by single-cell RNA sequencing. Functionally, GDF15^high^ macrophages exhibit dampened responsiveness to pro-inflammatory activation and can also inhibit the inflammatory responses of other macrophages via a paracrine mechanism. GDF15 *per se* acts as at least partly the anti-inflammatory mediator released by GDF15^high^ macrophages. Moreover, GDF15^high^ macrophages are found to be present in multiple human tissues in the body. These data suggest that GDF15^high^ macrophages may represent a novel macrophage population with intrinsic anti-inflammatory functions, although their (patho)physiological importance remains to be explored.

## Acknowledgments

This study was partially supported by research grants from National Natural Science Foundation of China (82070265 for F.J.; 82070820 and 82170495 for Q.Z.), Shandong Provincial Natural Science Foundation (ZR2021MH111 for X.C.), National Science and Technology of China (2012ZX09303016-003 for H.G.), and National Key R&D Program of China (2021YFA1301102 for Q.Z.).

## Author contributions

C.D. was involved in sample preparation, experimentation, data acquisition and data analysis; H.Z. was involved in sample preparation, data acquisition and data analysis; Z.Z. was involved in scRNA-seq data processing, analysis, visualization and bioinformatics mining; C.G.L. was involved in result interpretation and manuscript writing; M.M. was involved in experimentation and data acquisition; H.G. was involved in study design, result interpretation, and project management; Q.Z. was involved in study design, scRNA-seq data analysis and result interpretation; X.C. was involved in study conception, study design, experimentation, data analysis, and result interpretation; F.J. was involved in study conception, study design, result interpretation, and manuscript writing.

