## Supplementary Figures and Tables for "Identification of a distinct cluster of GDF15^high^ macrophages exhibiting anti-inflammatory activities"

### Supplementary Figure S1

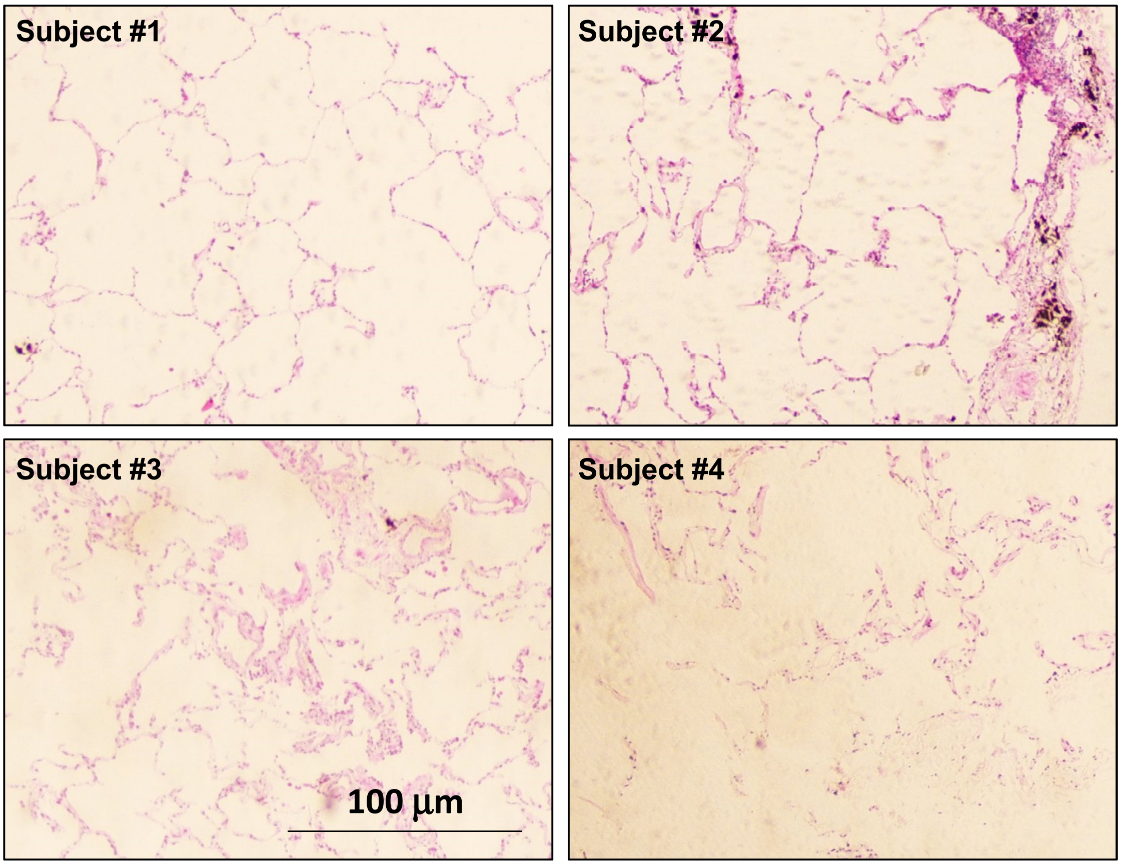


Figure S1. General histopathology of lung tissues from 4 individual patients with COPD. H&E stained sections showed that Subjects #1 and #4 had severe emphysema, resulting in poor cellularity of the lung tissue; in comparison, the lungs of Subjects #2 and #3 had significant inflammatory infiltrations.

### Supplementary Figure S2

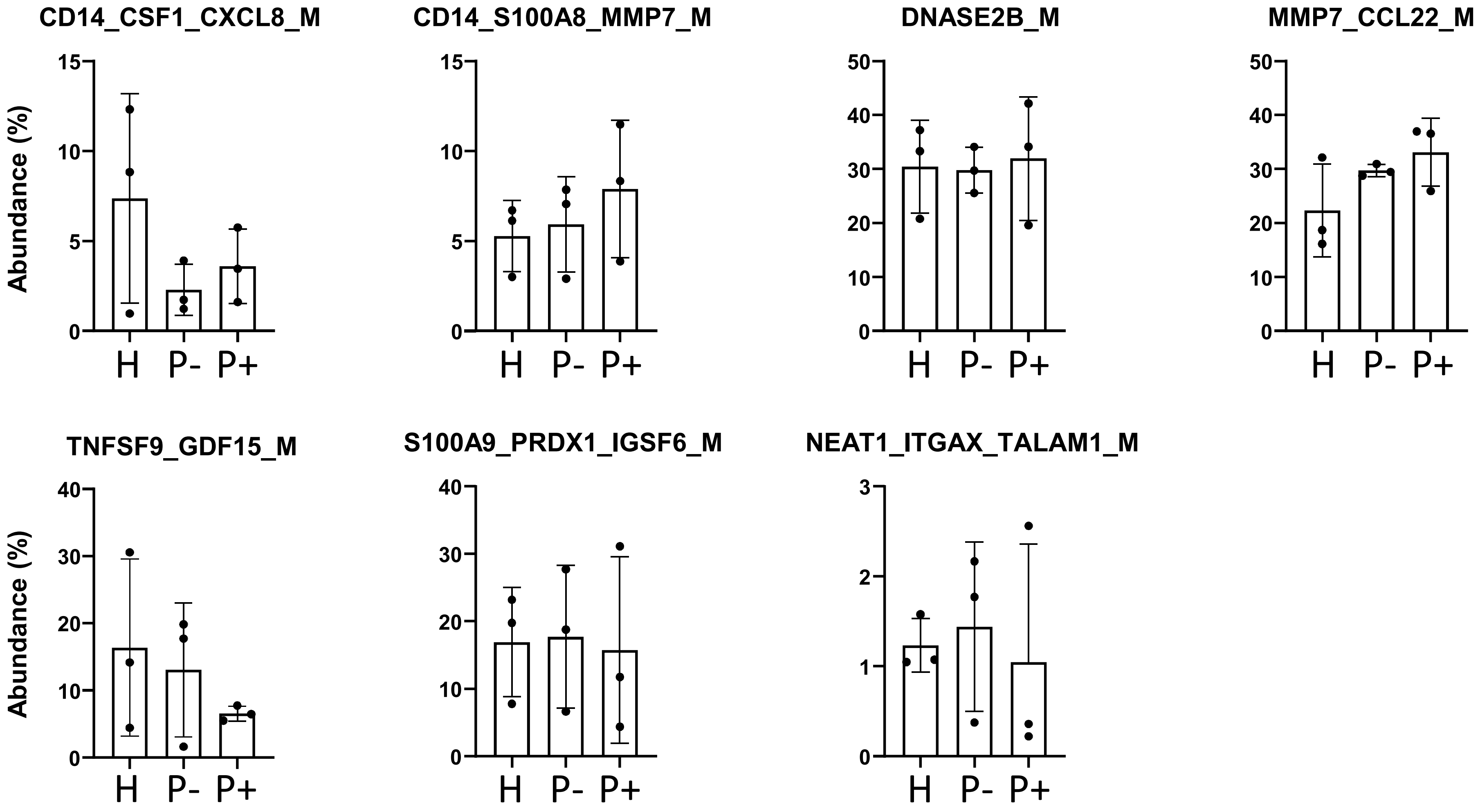


Figure S2. Quantitative data showing the percentage abundance of the 7 sub-populations of macrophages derived from human peripheral blood mononuclear cells. H, healthy subjects; P-, PAH patients without BMPR2 mutations; P+, PAH patients with BMPR2 mutations. Bars represent mean ± SD.

### Supplementary Figure S3

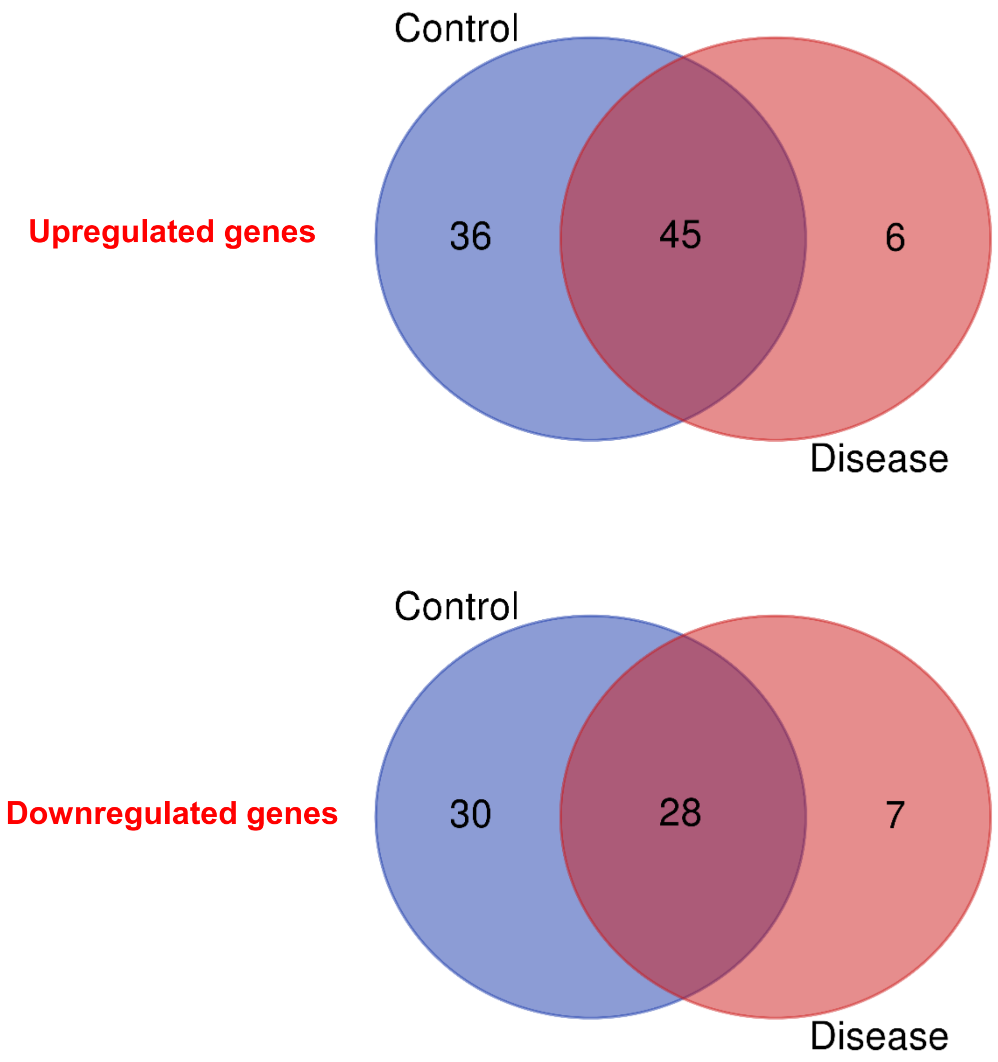


Figure S3. Venn diagrams showing the numbers of differentially expressed genes between GDF15$^{high}$ and GDF15$^{low}$ macrophages derived from 3 healthy subjects (Control) and 6 patients with pulmonary arterial hypertension (Disease), based on the scRNA-seq data set.

### Supplementary Figure S4

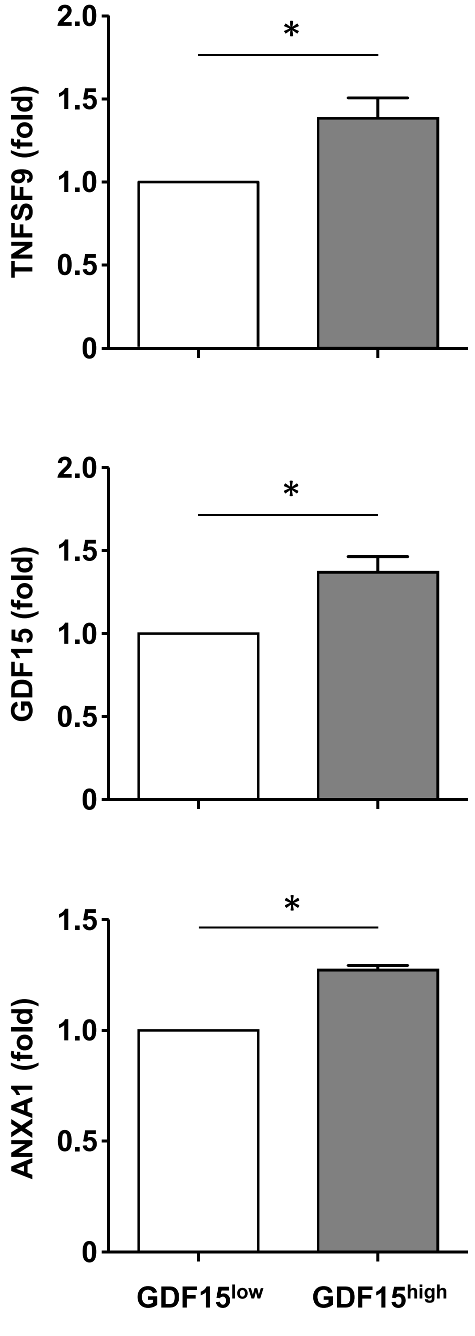


Figure S4. Results of ELISA assays for TNFSF9, GDF15 and annexin A1 (ANXA1) measured in the conditioned medium of rat BMMNC-derived macrophages. $GDF15^{low}$ and $GDF15^{high}$ cells were separated by FACS using TNFSF9 as a substitute marker. Data were expressed as mean ± SEM. * $P < 0.05$, unpaired *t*-test ($n = 3$ independent samples).

### Supplementary Figure S5

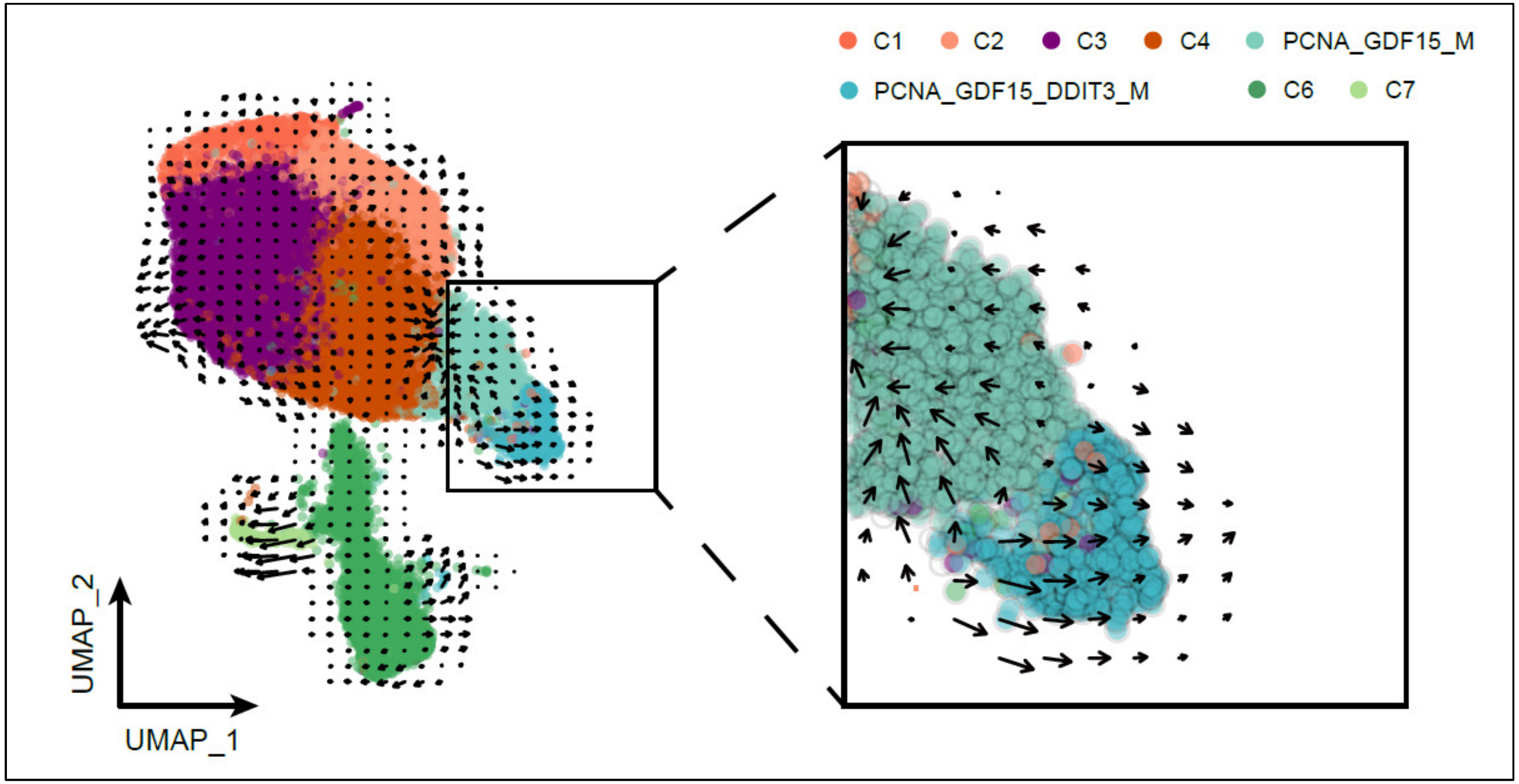


Figure S5. RNA velocity analysis result showing that the 2 sub-clusters of GDF15$^{high}$ macrophage might represent cells of the same identity but with diverging differentiation potentials; it was unlikely that these sub-clusters denoted two sequential differentiation status on the same differentiation route.

### Supplementary Figure S6

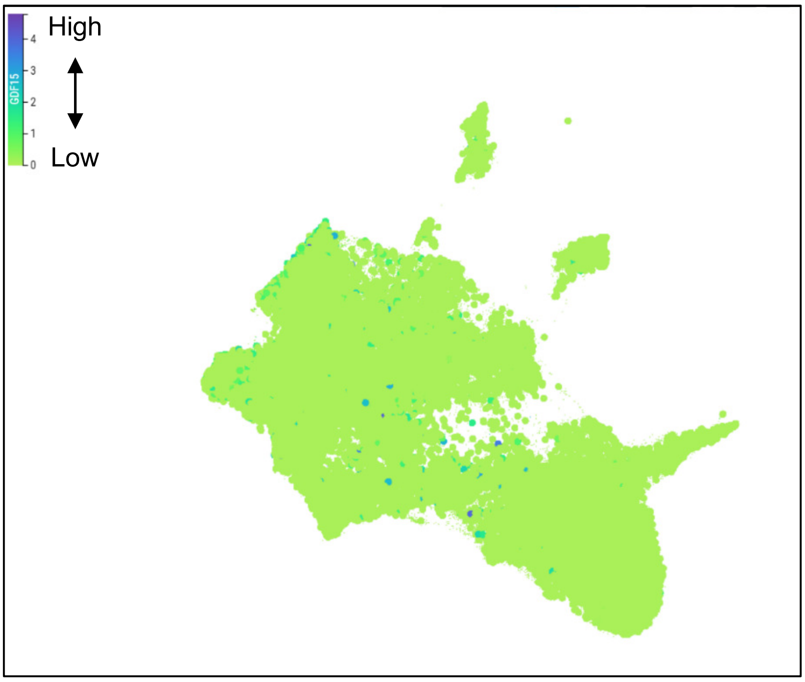


Figure S6. Analysis of the data in a public scRNA-seq database, MNP-VERSE, revealed that there was not a discernable cluster of monocytes with enhanced expression of GDF15. The color scale represented the relative expression level of GDF15.

**Supplementary Table S1. Basic demographic data of human subjects**

| | Total number | Gender | Age | Disease condition |
|---|---|---|---|---|
| **Healthy volunteers** | 7 | All females | 28-32 | N/A |
| **PAH patients** | 6 | All females | 28-32 | Three patients had bone morphogenetic protein receptor type 2 mutations; all patients were on ambrisentan and tadalafil therapies |

**Supplementary Table S2. Sequences of the primers used for real-time PCR**

| ***Rat*** | ***Forward*** | ***Reverse*** |
|---|---|---|
| β-actin | CTCTGTGTGGATTGGTGGCT | CGCAGCTCAGTAACAGTCCG |
| IL-1β | GGGATGATGACGACCTGCTA | ACAGCACGAGGCATTTTTGT |
| TNF-α | ATGGGCTCCCTCTCATCAGT | GCTTGGTGGTTTGCTACGAC |
| IL-6 | TTTCTCTCCGCAAGAGACTTCC | TGTGGGTGGTATCCTCTGTGA |
| ***Mouse*** | ***Forward*** | ***Reverse*** |
| β-actin | GGCTGTATTCCCCTCCATCG | CCAGTTGGTAACAATGCCATGT |
| IL-1β | GTGTCTTTCCCGTGGACCTT | AATGGGAACGTCACACACCA |
| TNF-α | CGGGCAGGTCTACTTTGGAG | ACCCTGAGCCATAATCCCCT |
| IL-6 | CTTCTTGGGACTGATGCTGGT | CTCTGTGAAGTCTCCTCTCCG |

**Supplementary Table S3. List of differentially expressed genes between GDF15high and GDF15low macrophages derived from 3 healthy subjects (Control) and 6 patients with pulmonary arterial hypertension (Disease)**

| Upregulated genes | | Downregulated genes | |
|---|---|---|---|
| ***Subjects*** | ***Gene symbol*** | ***Subjects*** | ***Gene symbol*** |
| Control & Disease | PCNA | Control & Disease | CYBB |
| | OMG | | ACTB |
| | MT1G | | LIMS1 |
| | DDIT3 | | ANXA2 |
| | MT1E | | APOC2 |
| | PMAIP1 | | COTL1 |
| | GAL | | TMSB4X |
| | ACTA2 | | NCAPH |
| | GCC2 | | HCST |
| | ZFAS1 | | C1QA |
| | PLIN2 | | ARPC1B |
| | TNFSF9 | | MNDA |
| | SNHG16 | | CHI3L1 |
| | CXCR4 | | TUBA1B |
| | ISCU | | TM4SF19 |
| | RRM2B | | ITGAX |
| | GADD45A | | FBP1 |
| | GAS5 | | IGSF6 |
| | TXNIP | | TMSB10 |
| | MT1M | | LGALS1 |
| | ANXA1 | | FDX1 |
| | GDF15 | | MSR1 |
| | RGS12 | | C1QB |
| | FDXR | | VIM |
| | ID3 | | FCER1G |
| | YBX3 | | ACTG1 |
| | MDM2 | | CTSB |
| | NCF1 | | ARHGDIB |
| | CXCL8 | Control only | ANXA5 |
| | HIST1H1C | | CTSD |
| | RPS27L | | TPM4 |
| | BTG1 | | HLA-DQB1 |
| | STK17A | | RPL37A |
| | PHLDA3 | | ATOX1 |
| | TP53I3 | | CYP1B1 |
| | ARPP19 | | SPARC |
| | CDKN1A | | ATP5F1E |
| | CD79A | | TGM2 |
| | RBP1 | | DBI |
| | MT1X | | CALM2 |
| | AK3 | | SMIM25 |
| | CRIPT | | FABP5 |

| | | | |
|---|---|---|---|
| | MT2A | | ITGB1BP1 |
| | MT1H | | TUBB |
| | SESN2 | | HAMP |
| Control only | SLC35E3 | | BCL2A1 |
| | SLC3A2 | | CTSZ |
| | CSRP2 | | CAPG |
| | HIST1H2AC | | RASSF4 |
| | METTL9 | | FCGR1A |
| | RRAD | | GALM |
| | TNFAIP6 | | TUBA1A |
| | TIMP3 | | S100A9 |
| | HIST1H2BJ | | ZFYVE16 |
| | MAP4K4 | | CIR1 |
| | UPP1 | | GAPDH |
| | BNIP3 | | C1orf162 |
| | ABCG1 | | SH3BGRL3 |
| | NEAT1 | Disease only | HLA-DRB1 |
| | CLU | | AC020656.1 |
| | ADA | | GBP1 |
| | C12orf49 | | SPP1 |
| | H2AFJ | | CCND1 |
| | C15orf48 | | STAT1 |
| | G0S2 | | ITGB2 |
| | YPEL3 | | |
| | DDB2 | | |
| | TAGLN2 | | |
| | KLK4 | | |
| | ST13 | | |
| | MLF2 | | |
| | SNHG32 | | |
| | PKD2L1 | | |
| | SLAMF7 | | |
| | MT-CO1 | | |
| | DUSP14 | | |
| | S100A10 | | |
| | CHI3L2 | | |
| | APOE | | |
| | FAS | | |
| | PTGDS | | |
| Disease only | NMB | | |
| | RPS19 | | |
| | AC006967.3 | | |
| | NMRK2 | | |
| | EIF1B | | |
| | CHIT1 | | |
